# miR-644a is a tumor cell-intrinsic mediator of sex bias in glioblastoma

**DOI:** 10.1101/2024.03.11.584443

**Authors:** Ellen S. Hong, Sabrina Z. Wang, András K. Ponti, Nicole Hajdari, Juyeun Lee, Erin E. Mulkearns-Hubert, Josephine Volovetz, Kristen E. Kay, Justin D. Lathia, Andrew Dhawan

## Abstract

**Background:** Biological sex is an important risk factor for glioblastoma (GBM), with males having a higher incidence and poorer prognosis. The mechanisms for this sex bias are thought to be both tumor intrinsic and tumor extrinsic. MicroRNAs (miRNAs), key post-transcriptional regulators of gene expression, have been previously linked to sex differences in various cell types and diseases, but their role in the sex bias of GBM remains unknown.

**Methods:** We leveraged previously published paired miRNA and mRNA sequencing of 39 GBM patients (22 male, 17 female) to identify sex-biased miRNAs. We further interrogated a separate single-cell RNA sequencing dataset of 110 GBM patients to examine whether differences in miRNA target gene expression were tumor cell intrinsic or tumor cell extrinsic. Results were validated in a panel of patient-derived cell models.

**Results:** We identified 10 sex-biased miRNAs (*a_djusted_ <* 0.1), of which 3 were more highly expressed in males and 7 more highly expressed in females. Of these, miR-644a was higher in females, and increased expression of miR-644a target genes was significantly associated with decreased overall survival (HR 1.3, *p* = 0.02). Furthermore, analysis of an independent single-cell RNA sequencing dataset confirmed sex-specific expression of miR-644a target genes in tumor cells (*p* < 10^-15^). Among patient derived models, miR-644a was expressed a median of 4.8-fold higher in females compared to males.

**Conclusions:** Our findings implicate miR-644a as a candidate tumor cell-intrinsic regulator of sex-biased gene expression in GBM.

**Key Points:**

- miR-644a is more highly expressed in female GBM patients.
- Lower miR-644a target gene expression is associated with improved overall survival.
- miR-644a target genes are higher in male GBM cells but not in other cell types.

**Importance of the Study:** MicroRNAs (miRNAs) are non-coding RNAs that regulate gene expression at the post-transcriptional level and were previously linked to glioblastoma (GBM) growth and therapeutic resistance. miRNAs play a role in the sex bias of various cell types and diseases, but how miRNAs contribute to sex differences in GBM is not well elucidated. We show that 10 miRNAs are differentially expressed between males and females and identify miR-644a as more highly expressed in female GBM patients. Using single-cell RNA-seq data, we demonstrate that sex differences in miR-644a target gene expression are tumor cell-intrinsic. Likewise, decreased miR-644a target gene expression is associated with improved overall patient survival. Our findings reveal miR-644a as a novel sex-biased miRNA in GBM, and a possible target for sex-specific precision therapies with limited collateral damage.

## Introduction

Glioblastoma (GBM) is the most common primary malignant brain tumor in adults. Despite an increasing scientific understanding of disease biology^1,2^, the median overall survival of patients with GBM is 17 months and has not improved since 2005^3^. The intrinsic resistance of tumor cells to therapy, as mediated by mechanism such as transcriptional programs and an immune-suppressive microenvironment, is the subject of significant investigation^4,5^.

Male biological sex has emerged as a key risk factor for both increased incidence and decreased survival in GBM^6,7^. This sex bias is believed to arise from both tumor-intrinsic and tumor-extrinsic factors. Tumor-intrinsic sex differences include differences in enhancer landscapes, glutamine metabolism, and oncogenic transformation^8,9^. Tumor-extrinsic factors contributing to sex bias include microglial activation and anti-tumor immunity^10,11^. However, the mechanisms underlying these sex-specific differences remain unknown.

We investigated microRNAs (miRNAs) as possible mediators of tumor-intrinsic sex differences in GBM. miRNAs are 19- to 25-nucleotide single-stranded RNAs that negatively regulate the expression of target genes through base pairing to the 3’ untranslated region of the mRNA transcript^12^. miRNAs are important for tumorigenesis, globally dysregulating the transcriptome in numerous cancer types^13^. In GBM, miRNAs, such as miR-7, have been shown to play key roles in tumor growth by repressing transcripts such as *EGFR*^14^. Furthermore, miRNA profiles of tumors have been shown to reliably subtype GBM and predict prognosis^15,16^, highlighting their clinical utility.

The role of miRNAs is context and cell type dependent, highlighting the need for miRNAs to be evaluated in specific cell types and diseases^17,18^. Notably, over 100 miRNAs are encoded on the X chromosome, while only 2 miRNAs are encoded on the Y chromosome^19^, which has led to studies of the association between miRNAs and sex differences. Additionally, both male and female sex hormones are known to influence miRNA expression^20,21^. Sex differences in miRNA expression have been reported in multiple tissues, including lung and brain^22,23^. Various diseases including lupus and metabolic syndrome also display a sex bias in miRNA expression^24,25^.

Sex bias in miRNA expression has also been reported in multiple cancers. These cancer types include those that are sex-specific (prostate, uterine) or have a substantial sex bias in terms of incidence, such as breast^26,27^. However, cancers with a less pronounced sex bias, such as colorectal cancer, lung adenocarcinoma, and hepatocellular carcinoma^28–30^, also exhibit a sex difference in miRNA expression. How miRNA expression differs between males and females with GBM and contributes to sex differences is unknown. Here, we leverage paired miRNA and mRNA sequencing of GBM samples to investigate sex-biased miRNA expression, and report in a tumor-intrinsic manner, the increased expression of miR-644a in biological females with associated repression of miR-644a target genes.

## Materials and Methods

### miRNA and mRNA analysis using patient Data from Kashani et al and GlioVis

We retrieved data from the Kashani et al^31^ and GlioVis^32^, which utilizes data from the Cancer Genome Atlas (TCGA)^33^. Briefly, Kashani and colleagues performed RNA sequencing on 100 ng total RNA using the nCounter_Human_miRNA_Expression_Panel_Assay_Kit H_miRNA_V3 or the nCounter_Human_PanCancer_pathway_panel (NanoString, Seattle, USA) spiked with 30 additional genes implicated in autophagy, epithelial mesenchymal transition, and DNA repair processes.

For the TCGA dataset, miR-644a target gene expression was obtained for the “TCGA GBM” cohort through the GlioVis web browser^32^. Additionally, matching clinical data for patients in the “TCGA_GBM” cohort was downloaded.

### Differential expression analysis

Differential expression analysis of the miRNA sequencing was performed using DESeq2 (v1.30.1)^34^. miRNAs with an adjusted p-value <0.1 were selected as differentially expressed miRNAs. R v4.0.5 was used.

### miRNA gene target prediction

miRNA gene targets were predicted using miRNAtap (v1.24.0)^35^. The default settings of including all 5 possible target databases were used: DIANA version 5.0, Miranda 2010 release, PicTar 2005 release, TargetScan 7.1, and miRDB 5.0. The default minimum source number of 2 was used, and the union of all targets found was taken as the initial set of targets for a given miRNA. miR-644a target genes were further filtered by removing target genes whose expression had positive Spearman’s correlations with miR-644a expression.

### Gene Ontology analysis

Gene ontology (GO) analysis was performed using www.geneontology.org. The default settings of searching for “biological process” with species “*Homo sapiens*” were used.

### Kaplan Meier curve

The Kaplan Meier survival plot was generated using R package survminer (v0.4.9). The fit was performed on survival, in days, and biological sex of patient. The log rank test was used for significance calculations.

### Survival analysis

The Cox proportional hazard ratio was calculated using the R package survival (v3.2.10). The miR-644a target gene expression score was defined as the mean of the log-normalized expression of negatively correlated miR-644a target genes. The variables included in the Cox model for overall survival include: miR-644a target gene expression score, biological sex, and age. miR-644a target gene expression score was calculated by taking the mean expression of the genes *MAKP9* and *PTPRR*.

### Single-cell RNA-sequencing data and analyses

The “Core GBmap” was downloaded from https://cellxgene.cziscience.com/collections/999f2a15-3d7e-440b-96ae-2c806799c08c^36^. Seurat (v4.3.0) was used to process the data. Default Seurat pre-processing steps were used, including: NormalizeData, FindVariableFeatures, ScaleData, RunPCA, FindNeighbors, FindClusters. Only cells that originated from a patient with known biological sex were kept. Only cell types that had at least 500 cells per sex were kept.

As above, miRNAtap was used to predict miR-644a targets. miR-644a target gene expression for each cell was retrieved, all genes with zero counts were removed from the analyses, and the mean expression of males and females was calculated across each cell type. The Wilcox rank-sum test was used to test for statistically significant differences between target expression in males and females on a cell-type specific basis.

To calculate z-scores for male to female miRNA target gene expression ratios, only genes that were expressed in at least 50% of all cells were retained. Next, the average log-normalized expression across all male and female cells for each miRNA and corresponding target gene was calculated. All above analyses were done using R v4.2.2.

### Real-Time Quantitative PCR

RNA was extracted from cells following the standard RNeasy kit protocol (Qiagen; 74134), and concentrations were measured using a NanoDrop (ThermoFisher) spectrophotometer. cDNA was synthesized using qScript cDNA SuperMix (Quanta Biosciences; 101414-102). qPCR was performed using Fast SYBR Green Mastermix (Applied Biosystems; 01120793) and an Applied Biosystems QuantStudio 3. Primer sequences are available in **Supplementary Table III**. During qPCR analysis, threshold cycle values were normalized to GAPDH.

### RNA-sequencing analysis

Expression of miR-644a targets in patient-derived GBM models was investigated using publicly available RNA-sequencing datasets. The L0, L1, L2 data was retrieved from E-MTAB-13161^37^. The HW1, PB1, RN1, SB2b, WK1 data was retrieved from Supplemental Dataset 4^38^. The 23M and DI318 data was retrieved from GSE119834^39^. Each individual model was rank-normalized, and the top 10% of miR-644a gene targets were evaluated. The mean of the ranks for miR-644a gene targets was calculated per model, and a one-sided Wilcox test was performed.

### Patient-derived cell models

L0, L1, and L2 were provided by Dr. Brent Reynolds (University of Florida) and have been described previously^40^. DI318 cells were obtained from the Rose Ella Burkhardt Brain Tumor Center biorepository (Cleveland Clinic) and have been described previously^41^. 3691 and 3832 were received from Dr. Jeremy Rich (University of Pittsburgh) and have been described previously^42^. 23M was received from Dr. Erik Sulman (New York University) and has been described previously^43^. SB2b, WK1, PB1, RN1, MMK1, and HW1 were received from Dr. Andrew Boyd (University of Queensland) and have been described previously^38^.

### Cell culture

All cells were grown at 37°C with 5% CO_2_. The L0, L1, L2, DI318, G3691, G3832, and GSC23 lines were grown in Neurobasal medium minus phenol red (Gibco) with 1X B-27 supplement (Gibco), 1 mmol/L sodium pyruvate, 2 mmol/L L-glutamine, 50 U/mL penicillin/streptomycin, 20 ng/ml hEGF and 20 ng/ml hFGF2 (R&D Systems). The remaining cell lines were grown in DMEM-F12 (Cleveland Clinic Media Preparation Core), 50 U/mL penicillin/streptomycin, 1% N2 supplement (Thermo Fisher Scientific), 20 ng/ml EGF and 20ng/ml FGF-2 (R&D Systems). Cells were passaged regularly using Accutase (Stem Cell Technologies) and phosphate-buffered saline.

### Data and Code Availability

All raw genomic data that were the source of analyses performed in this study have already been published. The matched miRNA and mRNA sequencing data were obtained from the authors of Kashani et al. 2023. The TCGA data obtained from GlioVis are available at http://gliovis.bioinfo.cnio.es/. The single-cell RNA-seq Seurat object is available at https://cellxgene.cziscience.com/collections/999f2a15-3d7e-440b-96ae-2c806799c08c. The RNA-sequencing data are previously published at E-MTAB-13161, GSE119834. All code necessary to reproduce the analyses in the figures of this manuscript is available on GitHub at https://github.com/esh81/miR644a_sexbias_GBM.

## Results

### Combined sample characteristics

For our comparative analysis, we leveraged tumor specimens from a cohort of 43 glioblastoma patients from the University of Bern during the years 1999–2016^31^. Importantly, this dataset contained matched mRNA and miRNA NanoString sequencing on tissue specimens from this cohort. The cohort consisted of 39 *IDH* wild-type patients, of whom 22 were biological males (herein referred to males), and 17 were biological females (herein referred to females) (**Supplementary Table I**). The median age at diagnosis was 57 years (range 41 to 79 years) (**Supplementary Table I**).

To validate the findings from the Kashani et al. cohort in a larger and independent cohort, the Cancer Genome Atlas (TCGA)^33^ database was utilized. Of the 538 patients, a total of 142 were *IDH* wild-type patients, with 93 male patients and 49 female patients (**Supplementary Table I**). The median age at diagnosis was 62 years (range 24 to 89 years) (**Supplementary Table I**).

### miR-644a shows significantly higher expression in GBM samples from females

Differential expression analysis (**Figure 1**) of the 22 females and 17 males in the Kashani et al. dataset for miRNA expression revealed statistically significant differences in 10 of 809 miRNAs (**Figure 2A**). Of these 10 microRNA, 3 microRNAs were more highly expressed in males and 7 were more highly expressed in females. While the fold changes are relatively modest between samples, this effect was not driven by outliers. A literature search suggested miR-644a as a candidate sex-biased miRNA in prostate cancer^44^, and we therefore examined this miRNA further.

**Figure 1.**
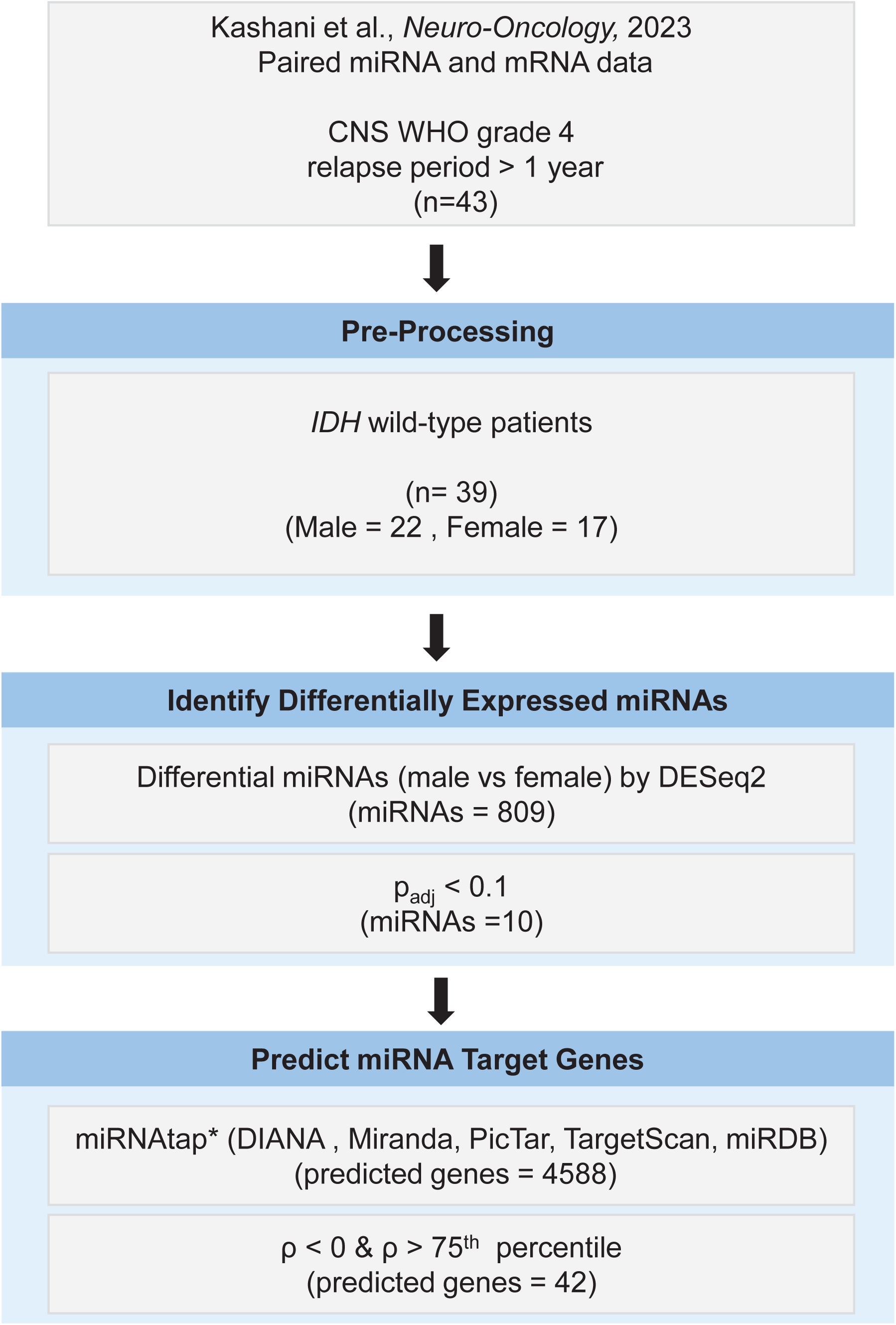
Analysis workflow to identify miRNAs differentially expressed in males versus females in *IDH* WT GBM. CNS (Central Nervous System). WHO (World Health Organization). *IDH* (isocitrate dehydrogenase). miRNA (microRNA). p_adj_ (p value adjusted). * = gene was considered a predicted target if at least 2 sources agreed.

**Figure 2.**
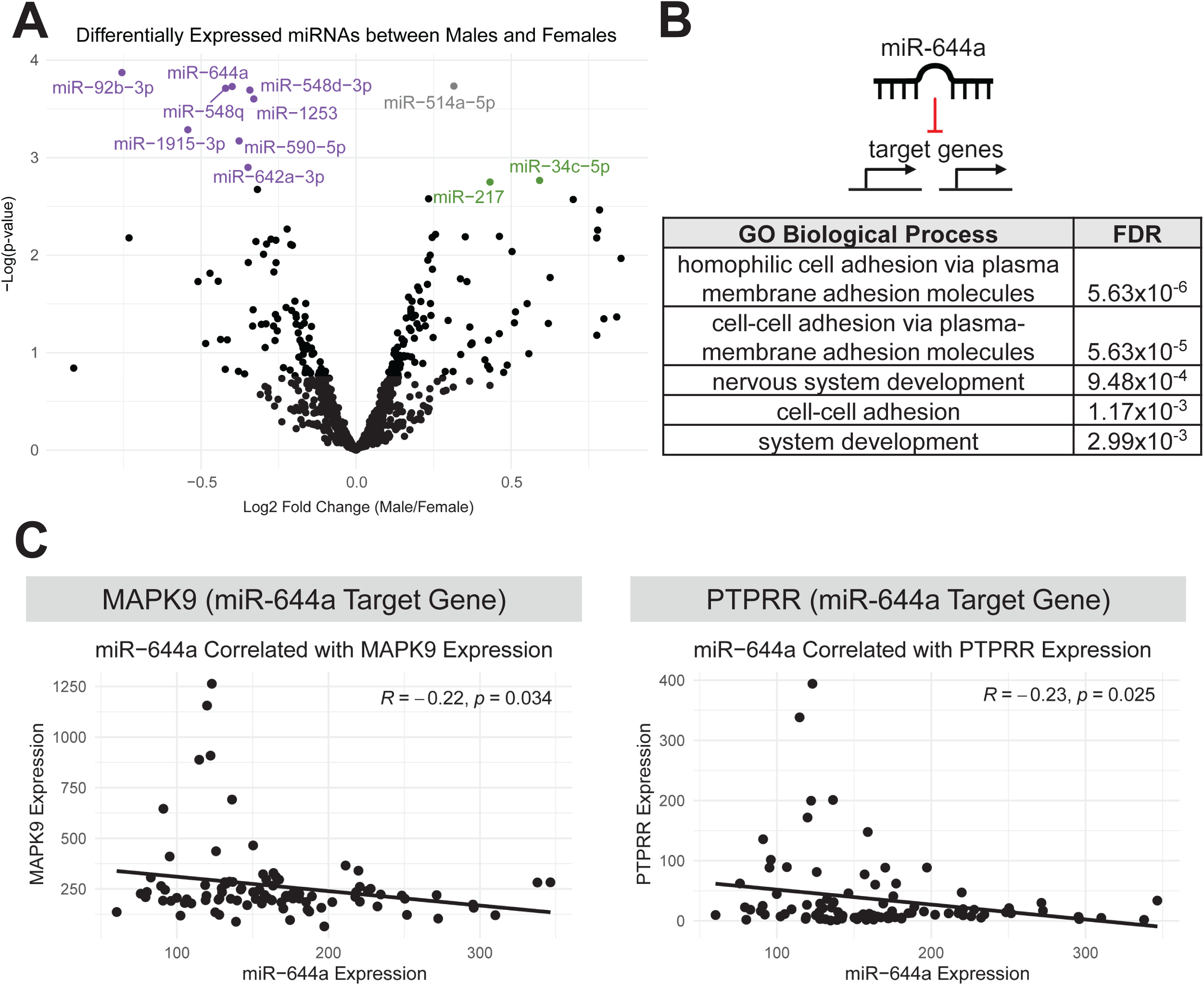
Differential expression analysis reveals sex-biased microRNAs. **A)** Volcano plot showing results of DESeq2 analysis. miR-514a-5p was excluded from further analyses due to its genomic location on the X chromosome. **B)** Top Gene Ontology (GO) Biological Processes for target genes predicted to be regulated by miR-644a. FDR (False Discovery Rate). **C)** Scatter plot of miR-644a expression in Kashani et al. dataset correlated with miR-644a target gene expression. (Left) MAPK9 and (right) PTPRR*. R =* Pearson’s correlation.

### miR-644a negatively regulates genes associated with cell adhesion and differentiation in GBM

We performed Gene Ontology analysis on predicted miR-644a targets, highlighting its regulation of genes known to act in cell adhesion and development (**Figure 2B**). Among the predicted mRNA targets negatively correlated with miR-644a expression (**Methods**, **Supplementary Table II**), *MAPK9* and *PTPRR* emerged as the most significant candidate target genes negatively regulated by miR-644a (**Figure 2C**). *MAPK9* is a kinase involved in the JNK signaling pathway and is associated with prognosis in GBM^45^, and *PTPRR* is a protein tyrosine phosphatase shown to inhibit *ERK1/2* in prostate cancer^46^. For all downstream analyses, *MAPK9* and *PTPRR* were used as miR-644a target genes in GBM.

### Increased miR-644a target gene expression is associated with decreased overall survival

To determine if miR-644a is associated with outcome in GBM patients, we performed survival analyses. Firstly, in the Kashani et al. cohort, we showed that males survive shorter than females (male median survival = 30 months, female median survival = 36 months, HR 1.5 log rank, *p* < 0.0001) (**Figure 3A**). We then examined the distribution of miR-644a expression in males versus females and found that as expected from the differential miRNA expression analysis (**Figure 2A**), females express 1.3-fold higher miR-644a compared to males (**Figure 3B**). Altogether, males in the Kashani et al. cohort have decreased miR-644a expression and reduced overall survival compared to females.

**Figure 3.**
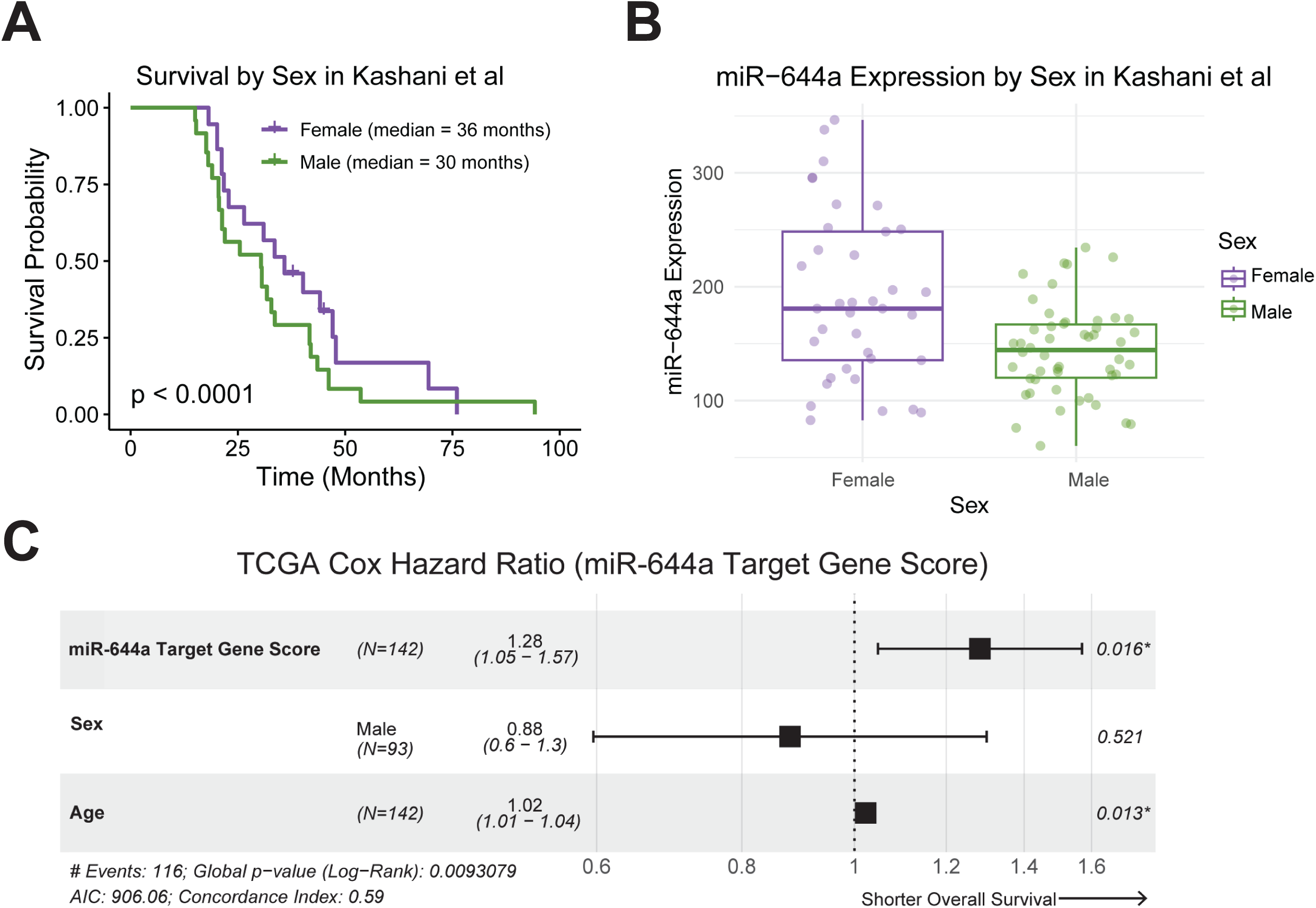
Increased miR-644a target gene expression is associated with decreased overall survival. **A)** Kaplan-Meier of overall survival in Kashani dataset. Log rank p-value. **B)** Boxplot of miR-644a expression by sex in Kashani et al. **C)** Cox proportional hazard ratio of miR-644a target gene score, sex, and age on overall survival of TCGA *IDH* wild-type glioblastomas.

To validate the analyses from the Kashani et al. cohort, we assessed the larger TCGA cohort, which has 142 patients. We performed a Cox proportional hazard ratio analysis on *IDH* wild-type tumors, evaluating survival as it related to miR-644a target gene expression, summarized into the miR-644a score, fully detailed in the Methods (**Figure 3C**). Multivariate Cox proportional hazards analysis on TCGA samples revealed an association between decreased overall survival and increased miR-644a target gene expression (HR 1.3 (95% CI 1.1 – 1.6), p = 0.02) (**Figure 3C**). In other words, higher expression of miR-644a target genes, which implies lower miR-644a activity, is associated with worse survival. As males have reduced expression of miR-644a and greater expression of miR-644a target genes, this finding suggests that reduced miR-644a expression in males may contribute to their poorer overall survival.

### miR-644a target genes are more highly expressed in male tumor cells compared to non-tumor cells

We then surveyed miR-644a target gene expression in an independent single-cell RNA-seq dataset to ascertain how miR-644a target gene expression varied among the cellular subtypes within GBM. We utilized the publicly available single-cell RNA-seq resource GBmap^36^, which curated over 300,000 single cells from 110 glioblastoma samples (27 females and 53 males). This resource annotated cells as one of 10 cell types, of which we retained 8 with sufficient representation across both males and females (**Figure 4A**).

**Figure 4.**
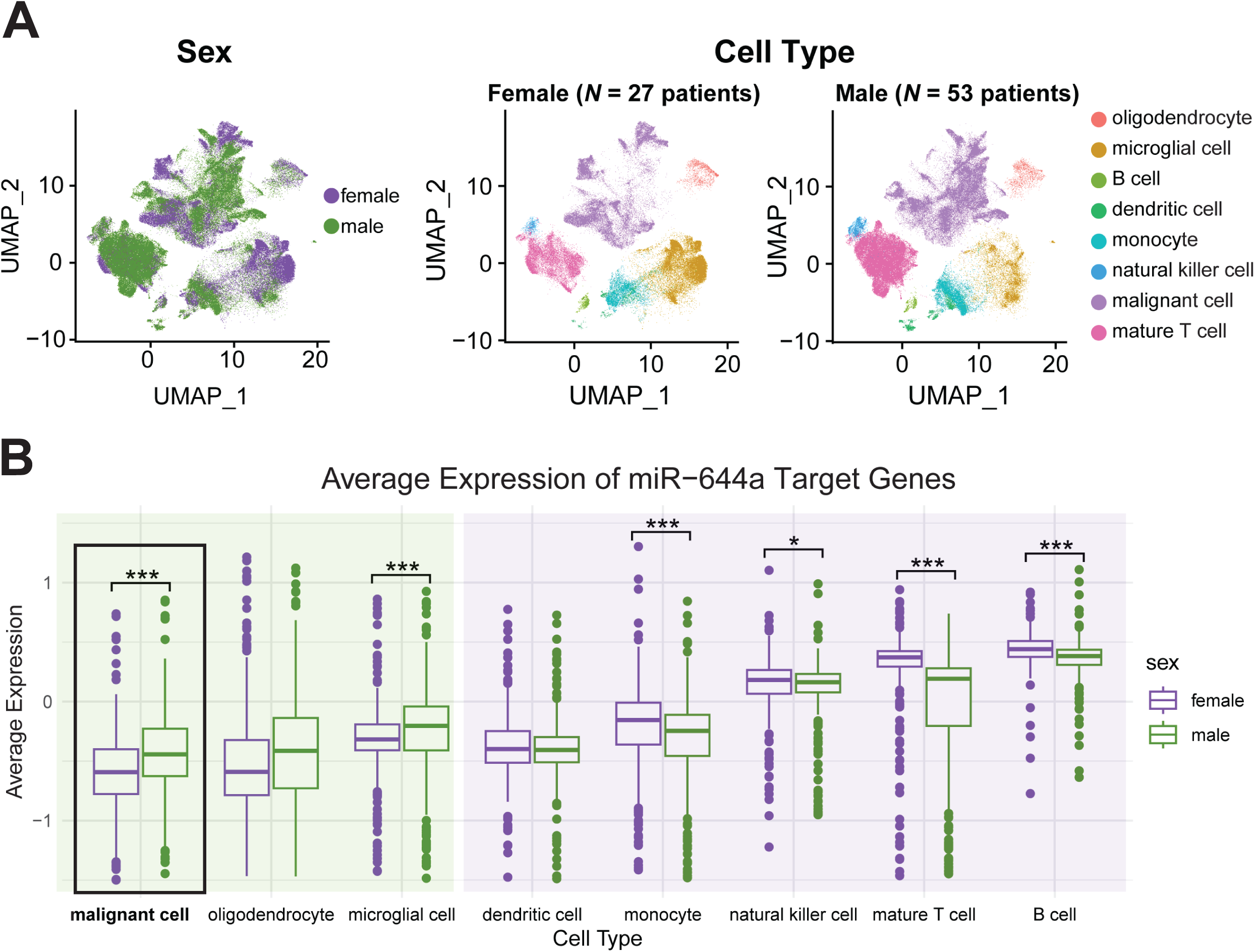
Male GBM cells more highly express miR-644a target genes. **A)** UMAP plots of sex (left) and cell type (right) from GBmap. **B)** Boxplot showing average expression (with zeros removed) of miR-644a target genes by cell type and sex in GBmap. Each point is one miR-644a target gene. ***p-value <0.001 and *p-value <0.05 using Wilcox test.

Because miRNA was not quantified in these patients, we used the average expression of miR-644a target genes as a surrogate for miRNA activity. At a single-cell level, we compared the average expression of miR-644a target genes in males versus females across each cell type (**Figure 4B**). Of all cell types, only malignant cells and resident brain cells, microglia, had significantly increased expression of miR-644a target genes in males (**Figure 4B**). In the remaining cell types, miR-644a target genes were more highly expressed in females compared to males. This suggests that miR-644a expression is driven in females by malignant cells, given the reduced expression of target genes in this cell subpopulation.

### miR-644a target genes are more highly expressed in male tumor cells relative to all other miRNAs

To elucidate if male tumor cells specifically express miR-644a target genes more highly, we inspected the landscape of all miRNA target genes across cell types. For each cell type, we predicted miRNA target genes for every known miRNA in the genome, and calculated the average expression in male and female cells (**Figure 5A**). We then normalized the distribution of male to female ratios for miRNA target genes per cell type (**Figure 5B**). Compared to target genes of all miRNAs, the z-score of miR-644a target gene expression for males over females was positive only among malignant cells and oligodendrocytes (**Figure 5C**), again suggesting the sex bias in expression of miR-644a target genes is specific to primarily tumor cells.

**Figure 5.**
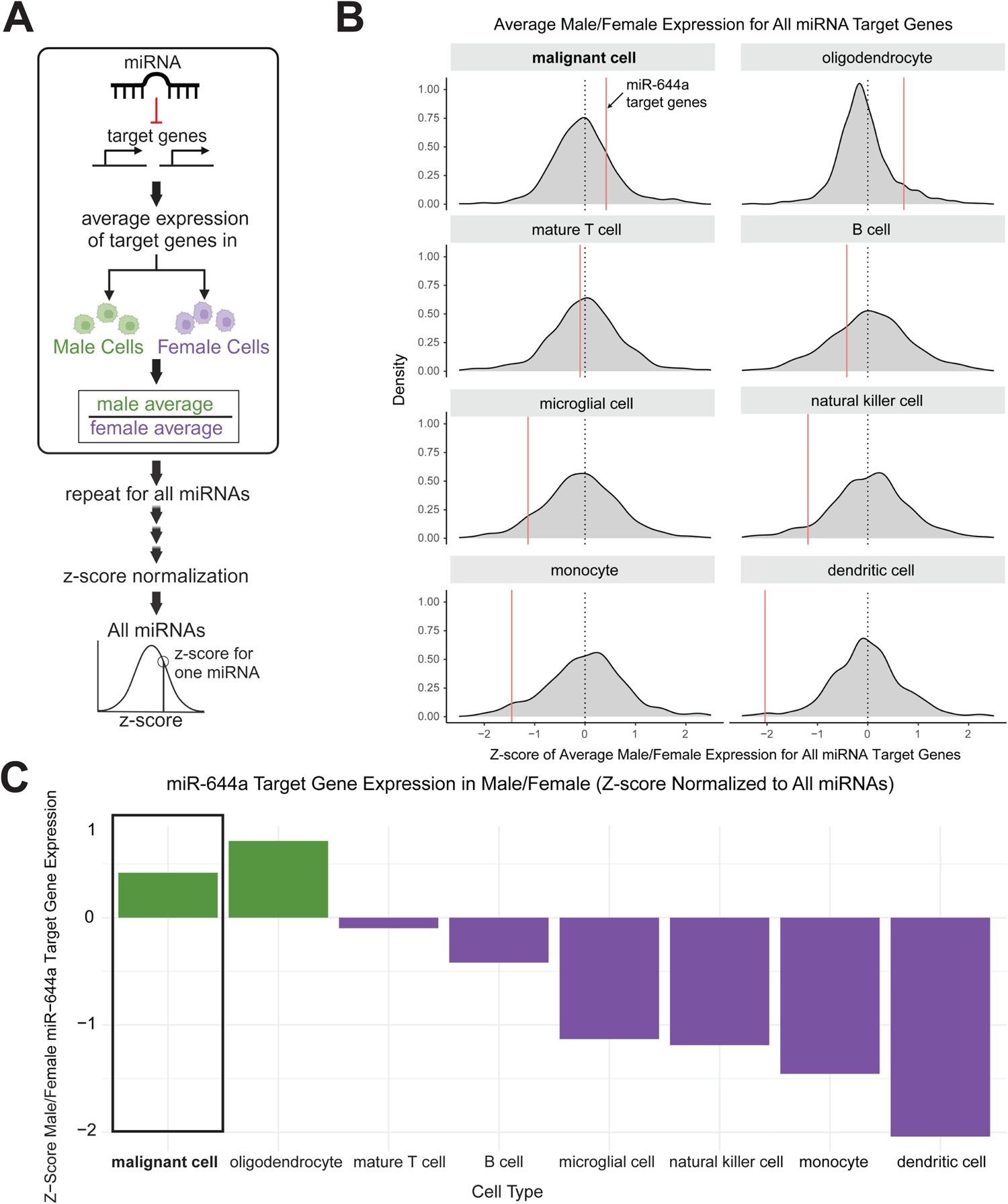
Male GBM cells more highly express miR-644a target genes relative to target genes of all other miRNAs. **A)** Schematic of z-score normalization for all miRNA target genes in all cell types. **B)** Z-score of distribution of male/female expression for all miRNAs. Red line shows average male/female expression for miR-644a target genes. **C)** Bar plot showing miR-644a target gene expression between males and females as a z-score within each cell type.

### miR-644a is more highly expressed in female patient-derived glioblastoma models

Finally, we validated miR-644a expression in our cohort of patient-derived glioblastoma models. We extracted total RNA from 12 total patient-derived glioblastoma models and assayed miR-644a expression. While there is heterogeneity across the different models, overall, female models express on average 4.8-fold more miR-644a than male models (**Figure 6A**).

**Figure 6.**
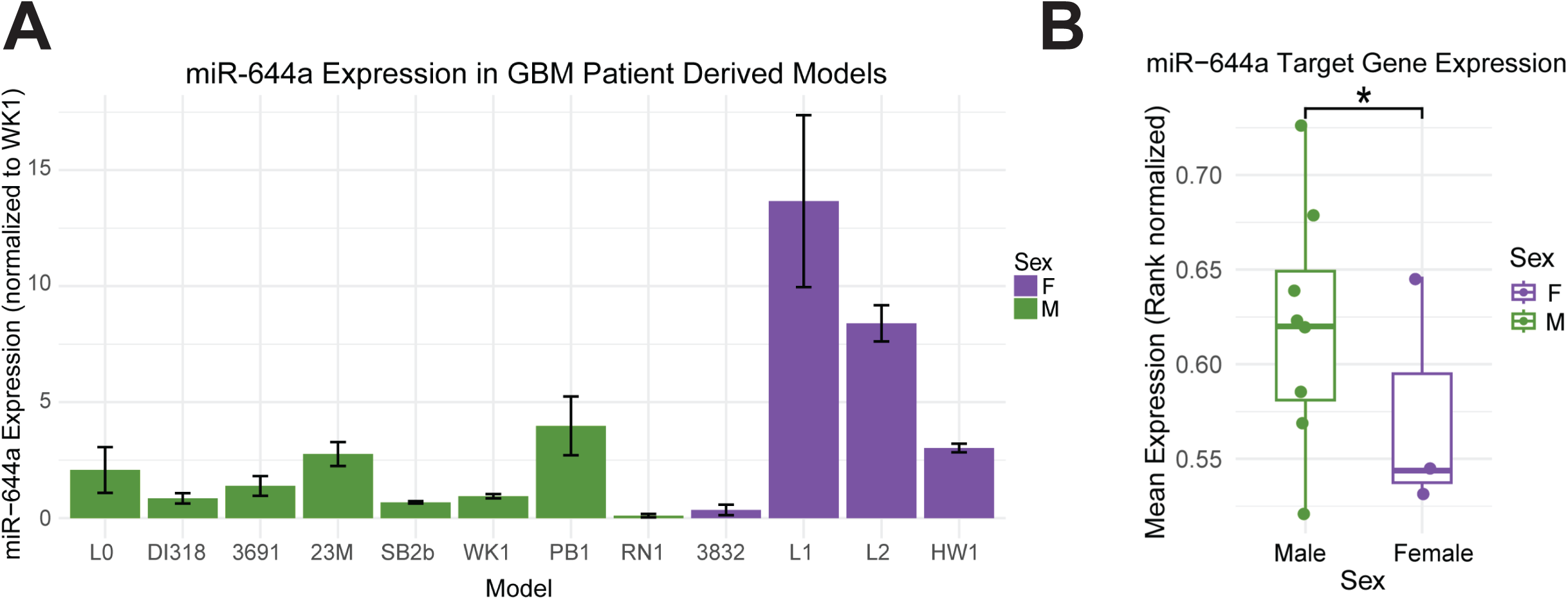
Female patient-derived GBM models more highly express miR-644a. qPCR of miR-644a in female and male patient-derived models. Every 1111C^T^ is the average of 3 biological replicates (normalized to *GAPDH*) and the fold change is normalized to WK1. **B)** Boxplot representing the average expression per model of miR-644a target genes (rank normalized per model). *One-sided Wilcox test *p*<0.05.

Moreover, we examined the expression of predicted miR-644a target genes in our patient-derived models and found, as expected, that females express miR-644a target genes at a lower level than males (*p* <0.05) (**Figure 6B**). These results illustrate that miR-644a is more highly expressed in female patient-derived models, which have corresponding lower miR-644a target gene expression.

## Discussion

miRNAs are key regulators of GBM growth, self-renewal, and therapeutic resistance. Here, we show that males and females with GBM differentially express a subset of miRNAs, and among these, miR-644a is more highly expressed in female patients in a tumor cell-intrinsic manner. Using the predicted target genes for miR-644a (*MAPK9* and *PTPRR*), we discover that higher expression of miR-644a target genes associates with decreased overall survival. Using single-cell RNA-sequencing data, we demonstrate that miR-644a target genes are also differentially expressed in male and female malignant cells compared to other cell types. Finally, we validate the higher expression of miR-644a in a panel of female GBM patient-derived models, relative to male patient-derived models. Taken together, our results across three independent datasets and 12 patient-derived models demonstrate that miR-644a is more highly expressed in females and may play a role in GBM sex bias.

Males have a higher incidence and poorer prognosis of GBM. However, the mechanisms of this sex bias are still an area of active investigation. miRNAs, as context-specific regulators of gene programs, warranted examination as a candidate mediator of sex-biased transcriptional differences. We demonstrated that increased expression of miR-644a target genes is associated with worse survival. The gene programs regulated by miR-644a are most prominently cell adhesion and development, which have also been linked to sex differences in GBM^47^. Therefore, it is possible increased miR-644a expression in females leads to decreased GBM progression compared to males. Furthermore, miR-644a target genes were significantly overlapped with previously reported male-specific genes but not female-specific genes^47^, supporting the function of miR-644a in repressing genes in females.

The mechanism behind the sex-biased expression of miR-644a remains unexplored. Unlike miRNAs encoded on the X chromosome that have been previously linked to sex differences, miR-644a is encoded on chromosome 20. Hence, the sex-biased expression of miR-644a is not entirely due to its chromosomal location. DNA methylation of miRNA promoters influences expression of the miRNA in various cancers such as colorectal cancer^48^. We therefore hypothesize that miR-644a may be differentially methylated in males versus females, contributing to its sex-specific expression, but this remains to be validated. Finally, aberrations in miRNA processing such as miRNA biogenesis and miRNA degradation can also affect miR-644a expression, but these have not yet been explored as contributors to sex differences in GBM^49,50^.

There are limitations to our current study. In general, there is a scarcity of work investigating miRNAs due to the challenges in assaying their expression and function, which limits the feasibility of studies. Due to the lack of datasets with matched miRNA and mRNA sequencing, we were restricted to a small sample size. While Kashani and colleagues performed matched miRNA and mRNA sequencing, the sequencing was a targeted panel and did not assay the full transcriptome. Furthermore, as there is no miRNA information at the single-cell level, miRNA expression had to be extrapolated by examining target gene expression. Additionally, as protein expression may not correlate with miRNA or mRNA levels, the functional impact for these findings is yet to be determined.

Overall, these findings contribute molecular complexity to sex bias in GBM. miRNAs are attractive therapeutic targets because they utilize the cell’s intrinsic machinery and act in context- and cell type-specific ways. We showed that miR-644a target genes were increased in males compared to females in malignant cells, which highlights the potentially limited collateral damage to other cell types in the tumor. Thus, in summary, we demonstrate that miR-644a is a mediator of sex differences in GBM, and a potential target for future tumor-intrinsic, sex-specific therapies.

## Acknowledgments

The authors thank all members of the Lathia and Dhawan laboratories for thoughtful discussion and project conceptualization.

## Funding

NIH grants R35 NS127083 (J.D.L) and P01 CA245705 (J.D.L). This work was also supported by the American Brain Tumor Association (J.L., J.D.L.), American Academy of Neurology grant AAN2313JS (A.D.), Case Comprehensive Cancer Center (J.D.L), and Cleveland Clinic/Lerner Research Institute (J.L., J.D.L, A.D.).

## Conflict of Interest

All authors declare no conflicts of interest.

## Authorship

E.S.H., J.D.L, and A.D. wrote the manuscript, with editing provided by J.L and E.E.M.H. E.S.H., J.D.L., and A.D. designed the study, and E.S.H. performed primary data collection and analysis. S.Z.W., J.L., E.E.M.H., J.V., and K.E.K. assisted with experimental design. S.Z.W., A.K.P, N.H. assisted with culture of patient-derived models and RNA extraction. S.Z.W., J.L., J.V., and K.E.K. assisted with interpretation of the data.

## Notes

### Competing Interest Statement

The authors have declared no competing interest.

