## Supplementary Materials for "miR-644a is a tumor cell-intrinsic mediator of sex bias in glioblastoma"

| | Kashani et al. (*N* = 39) | TCGA (*N* = 142) |
|---|---|---|
| *IDH* Wild-Type | 39 | 142 |
| Biological Sex | | |
| Male (%) | 22 (56%) | 93 (65%) |
| Female (%) | 17 (44%) | 49 (35%) |
| Median Age (years) | 57 | 62 |
| *MGMT* Methylation Status | | |
| Unmethylated (%) | 16 (41%) | 63 (44%) |
| Methylated (%) | 19 (49%) | 46 (33%) |
| No data (%) | 4 (10%) | 33 (23%) |

**Supplementary Table I. Summary of patient characteristics in datasets used.** TCGA (The Cancer Genome Atlas); *IDH* (isocitrate dehydrogenase); *MGMT* (O-6-Methylguanine-DNA Methyltransferase).

| gene | corr_coeff | p-value |
|---|---|---|
| ARNT2 | -0.153 | 0.157 |
| BMP4 | -0.012 | 0.911 |
| CAMK2B | -0.017 | 0.876 |
| CCR7 | 0.087 | 0.421 |
| CDK6 | 0.041 | 0.709 |
| CLCF1 | 0.103 | 0.344 |
| FGF11 | 0.136 | 0.211 |
| FGF7 | 0.160 | 0.139 |
| FLNC | 0.023 | 0.829 |
| MAPK9 | -0.213 | 0.048 |
| NF1 | -0.092 | 0.396 |
| NRAS | 0.059 | 0.586 |
| PAX3 | -0.036 | 0.742 |
| PITX2 | -0.061 | 0.575 |
| PRDM1 | -0.091 | 0.403 |
| PTPRR | -0.202 | 0.060 |
| TLX1 | 0.167 | 0.123 |
| TSPAN7 | 0.040 | 0.716 |

**Supplementary Table II. miR-644a gene targets.** miR-44a genes predicted by miRNAtap with Spearman’s correlation coefficients (corr_coeff) and p-values with miR-644a expression from Kashani et al. dataset.

| Primer | Sequence |
| --- | --- |
| miR-644a Forward | GCGGCGAGTGTGGCTTTC |
| miR-644a Reverse | CAGTGCAGGGTCCGAGGT |
| GAPDH Forward | GGAGCGAGATCCCTCCAAAAT |
| GAPDH Reverse | GGCTGTTGTCATACTTCTCATGG |

**Supplementary Table III. Primer sequences**.
